# Membrane tension propagation couples axon growth and collateral branching

**DOI:** 10.1101/2022.01.09.475560

**Authors:** Zheng Shi, Sarah Innes-Gold, Adam E. Cohen

## Abstract

Neuronal axons must navigate a mechanically heterogeneous environment to reach their targets, but the biophysical mechanisms coupling mechanosensation, growth, and branching are not fully understood. Here, we show that local changes in membrane tension propagate along axons at approximately 20 µm/s, more than 1000-fold faster than in other non-motile cells. This rapid and long-range mechanical signaling mediates bidirectional competition between axonal branch initiation and growth cone extension. Our data suggest a mechanism by which mechanical cues at one part of a growing axon can affect growth dynamics remotely.

## Introduction

Mechanical forces play an important role in modulating axonal growth and branching.^1^ Axons *in vivo* follow stiffness gradients toward softer regions.^2^ Very small forces (< 10 pN) exerted by magnetic nanoparticles can speed axon growth.^3^ Several studies have suggested the presence of long-range coordination between growth of different axon branches. In cultured rat hippocampal neurons, growth rates of distinct axonal branches are anti-correlated.^4^ In cultured locust neurons, anchoring of individual branches caused retraction of neighboring un-anchored branches.^5^ The signals coordinating these dynamics across long distances have not been identified.

Several lines of evidence suggest that membrane tension could play a role, at least locally, in axon dynamics. The mechanical pushing force of the growth cone is closely balanced with the local membrane tension, both having values of order 10 pN/μm.^6^ Membrane addition at the growth cone is essential for growth, and blocking membrane addition at the growth cone stops elongation.^7,8^ Membrane tension also appears to play a role in inhibiting collateral branch nucleation. Overcoming the tension barrier by mechanically pulling a lateral membrane tether from an axon can nucleate growth of new functional axonal branches.^9^

We thus sought to determine whether membrane tension was localized or dispersed in axons and growth cones; and if tension was dispersed, whether tension propagation mediated long-range coordination of axonal growth and branching. Recent measurements of plasma membrane tension propagation in non-motile cells found that local perturbations to tension remained localized for many minutes and that the membrane tension could be highly heterogeneous within a single cell.^10^ This absence of tension propagation was attributed to the flow resistance of membrane-cortex attachments (MCAs). Due to the long-range influence of obstacles on two-dimensional flows, a cell membrane punctuated by a high density of MCAs was shown to behave rheologically more like a gel than a fluid.^11–13^

In contrast, long-range tension propagation has been proposed to play a central role in coordinating the shape and motions of motile cells. For example, in neutrophils, actin polymerization at the leading edge is thought to cause global increases in membrane tension which suppress formation of secondary growth fronts.^14^ Similarly, in fish skin keratocytes, membrane tension is proposed to couple actin growth rates across the cell, giving these cells their characteristic shapes.^15–17^ A key assumption in these models is that membrane flows lead to rapid equilibrium of membrane tension throughout the cell so that the tension is approximately homogeneous,^18^ or has at most a slight gradient from front to back.^19^ However, tension propagation in motile cells has not been measured directly. There is a ∼10^6^-fold range in proposed tension propagation speeds between non-motile vs. motile cells (tens of minutes vs. milliseconds for a local tension perturbation to cross a cell). As a motile appendage of a non-motile cell, it is not *a priori* obvious how one should think about propagation of membrane tension in the growth cone and the axon.

The membranes of axons and growth cones are notably different from other cellular structures in several regards. A tension gradient arises between the growth cone and the soma which drives retrograde membrane flow.^20^ In chick sensory neurons, tether attachment points can easily slide along the axon, suggesting that the spacing between MCAs exceeds the tether diameter (which is typically 50 – 100 nm).^21^ In contrast, for tethers pulled on HeLa cells,^21^ neutrophils,^22^ fibroblasts, epithelial cells, and endothelial cells,^10^ the attachment point remains pinned, implying a much smaller spacing between MCAs than on axons. Lipids diffuse faster in distal axons compared to in regions near the soma.^23^ Additionally, the total MCA strength per unit area, was recently estimated to be much lower in axons than in fibroblasts, consistent with a lower density of MCAs in axons.^24^ Finally, local tension propagation has been proposed to mediate ultrafast coupling of vesicle fusion and nearby vesicle endocytosis in mouse hippocampal axons,^25^ and was recently shown to couple vesicle fusion and endocytosis in axon terminals of goldfish retinal bipolar neurons.^26^ Together, these results suggest that the plasma membrane may encounter an unusually low density of obstacles in the axon compared to in other cellular structures, and thus might support propagation of membrane tension.

## Results

### Membrane tension propagates along the axon but not dendrites

We probed membrane tension propagation using a dual-tether assay in axons of cultured rat hippocampal neurons (Fig. 1A, Methods). Using two micromanipulators, we pulled a pair of tethers from a single axon in a neuron expressing cytoplasmic eGFP (Methods, Supplementary Movie 1). We then dynamically stretched tether 1 and monitored the fluorescence in both tethers (Fig. 1B). The fluorescence reported the tether’s cross-sectional area, which is inversely proportional to the local membrane tension.^27^ Stretching tether 1 led to a clear response in tether 2, 40 µm away (Fig. 1C). This observation established that membrane tension propagated along the axon.

**Figure 1:**
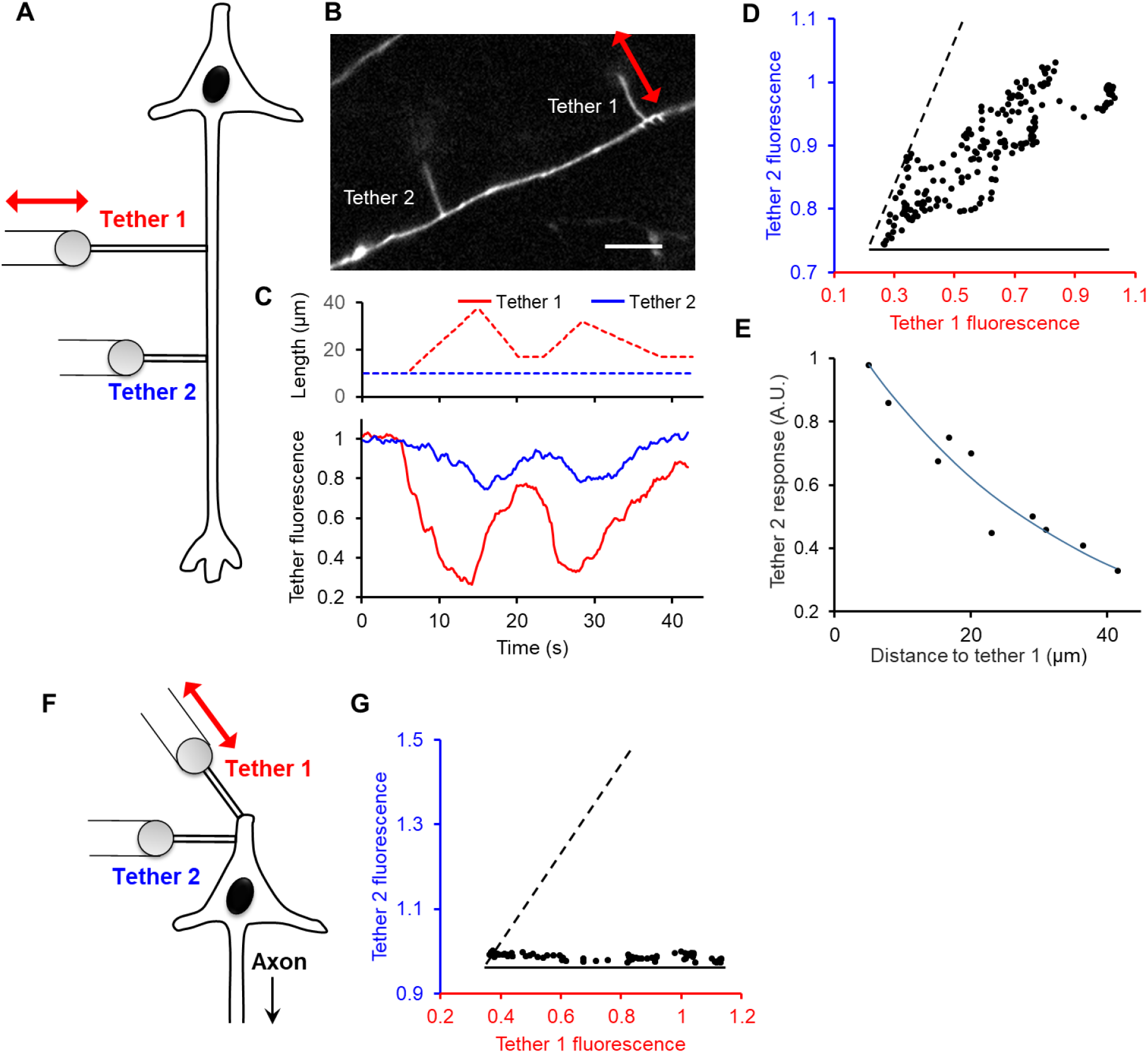
Membrane tension propagates in axons but not in dendrites. **A)** Double tether experiment to probe tension propagation in an axon. **B)** Two tethers, 40 µm apart, were pulled from an axon in a neuron expressing cytosolic eGFP. Scale bar 10 µm. **C)** Length (top) and fluorescence (bottom) of tether 1 (red) and tether 2 (blue) as tether 1 was stretched. **D)** Fluorescence of tether 2 as a function of fluorescence in tether 1. The fluorescence of tether 2 was time-shifted by 2 s to account for the tension propagation delay. Dashed and solid lines represent idealized responses for perfect tension coupling and zero coupling respectively. **E)** Ratio of fluorescence response in tether 2 to fluorescence response in tether 1 as a function of the separation between the tethers (*n* = 10 tethers, 10 neurons). Line is a fit to an exponential decay with length constant 32 μm. **F)** Double tether experiment to probe tension propagation on a proximal dendrite. **G)** Fluorescence of tether 2 as a function of fluorescence in tether 1. The two tethers were 15 µm apart. Dashed and solid lines represent idealized responses for perfect tension coupling and zero coupling respectively. Data are representative of experiments on *n* = 8 neurons.

We quantified the strength of the coupling as a function of distance between the tethers by measuring the ratio of the fluorescence changes in tether 2 to the changes in tether 1 for many tether pairs (*n* = 10 tether pairs, 10 neurons). The tension coupling was nearly perfect when the tethers were within 5 µm and decayed by half over approximately 30 μm (Fig. 1E). In contrast, no tension propagation was observed between two tethers pulled from proximal dendrites even when the tethers were less than 20 µm apart (Fig. 1F, 1G). The absence of tension propagation in dendrites is consistent with our earlier observations of no coupling between tethers in HeLa and other non-motile cells.^10^

### Axon pearls report membrane tension propagation speed

We found that tether-induced increases in local membrane tension could induce a pearling instability in the axon (Fig. 2, Supplementary Movie 2), similar to the pearling instability previously observed in axons under osmotic shock^28^ and in synthetic membrane tubes under high membrane tension.^29,30^ The pearling was reversible, disappearing when the tension was relaxed (Fig. 2C).

**Figure 2:**
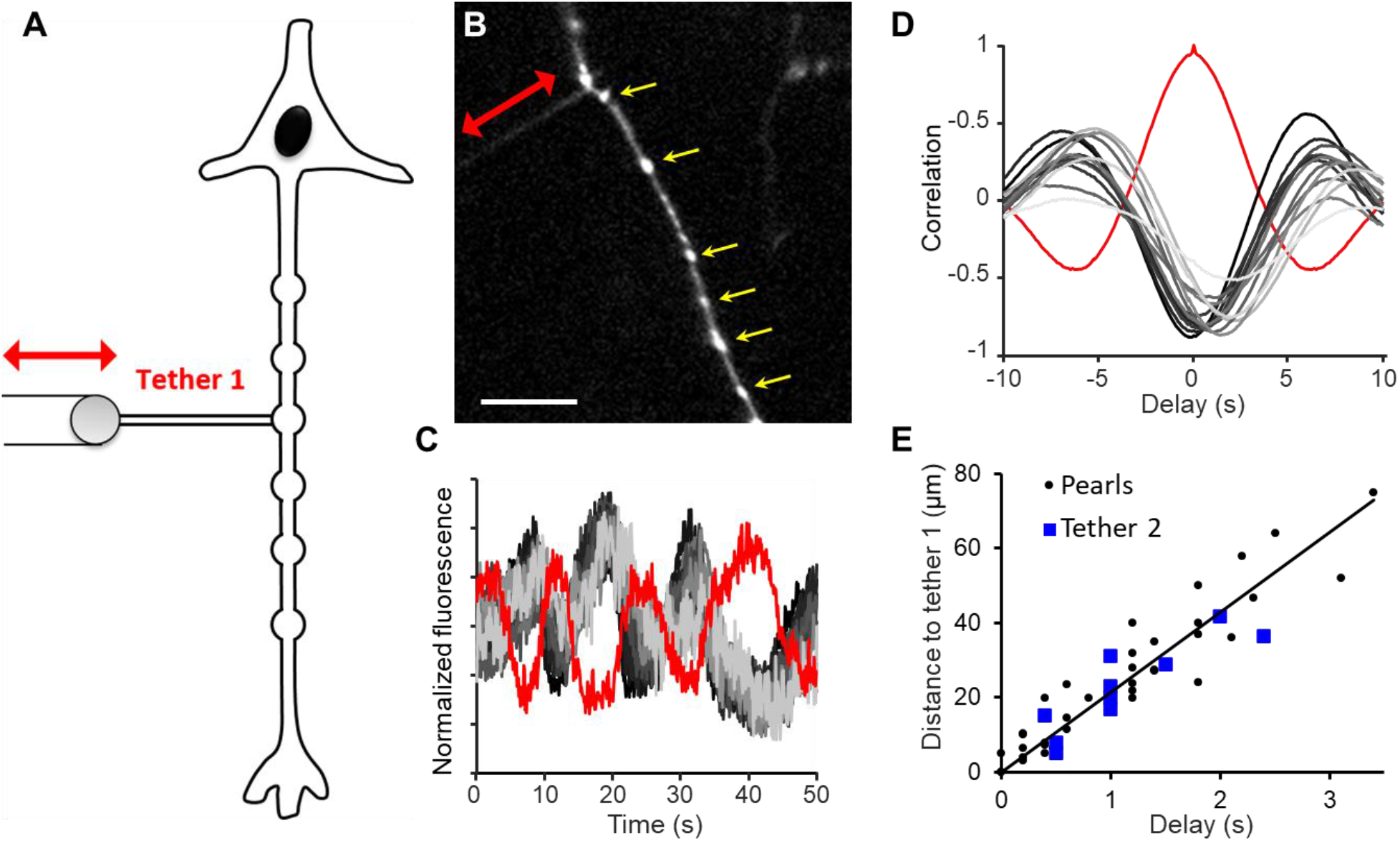
Membrane tension propagation triggers reversible pearling along the axon. **A)** Tension propagation speed can be measured by the distance-dependent time delay of tension-induced axon pearling. **B)** Image of an axon where pulling a tether induced pearls (yellow arrows). Scale bar 10 μm. **C)** Normalized fluorescence of tether 1 (red) and pearls (gray) on an axon of a neuron expressing cytosolic eGFP. Line shading goes from dark to light as pearl distance from tether increases (same in **D**). **D)** Red: autocorrelation of tether 1 fluorescence. Gray: cross-correlation between tether 1 fluorescence and pearl fluorescence. The time of the minimum near zero delay gives the time delay of each pearl. **E)** Delay for response of each pearl (black) or of a second tether (blue) as a function of distance to tether 1. A linear fit to the pearl data gives a velocity of 20.1 μm/s, *R*^2^ = 0.9.

Compared to the dual-tether assay, the pearling transition was a more convenient means to monitor tension propagation because pearling reported the tension dynamics all along the axon. We alternately stretched and relaxed single tethers and monitored the amplitude of the pearling as a function of distance and time (Fig. 2B, C). The pearling response lagged behind the tether stretch with a delay that increased with distance from the tether (Fig. 2D, Methods). The relation between time-lag and pearl distance gave a propagation speed of membrane tension of 22 ± 6 µm/s (mean ± s.d., *n* = 31 tethers, 30 neurons, Fig. 2E).

We validated this estimate of tension propagation speed using pairs of tethers pulled at a few discrete separations. The fluorescence of tether 2 responded to stretch of tether 1 with a lag that increased as a function of the separation, giving a similar tension propagation velocity to the pearl measurements (Fig. 2E).

### Sparse MCAs permit rapid tension propagation in axons

We next sought to determine why tension propagated so much further and faster in axons compared to all other cell types and structures we have studied.^10^ High mechanical stress at the growth cone has been shown to activate mechanosensitive ion channels, leading to a calcium influx which led to axonal retraction.^31^ We thus hypothesized that rapid diffusion of Ca^2+^ within the lumen of the axon might mediate changes in tension that followed the local Ca^2+^ dynamics.

We expressed axon-targeted GCaMP6s in cultured rat hippocampal neurons (Fig. S1).^32^ Breaking the axon led to a large increase in fluorescence, confirming proper expression and function of the Ca^2+^ indicator. However, we did not observe any Ca^2+^ signal during tether pulling. Even when tethers were pulled enough to elicit pearling, there was no observable Ca^2+^ signal (Fig. S1). These findings indicate membrane tension propagation could occur without Ca^2+^ influx.

Hydrodynamic models of membrane flow have shown that tension propagation is impeded by immobile MCAs,^10,11,33–36^ so we next sought to compare the density of MCAs in the axon to their density in dendrites. MCAs can obstruct diffusion of tracer molecules, though tracer diffusion is much less sensitive to MCA density than is tension propagation.^11,33^ To compare tracer diffusivity in different neuronal compartments, we performed fluorescence recovery after photobleaching (FRAP) experiments on rat hippocampal neurons expressing GPI anchored eGFP (Fig. S2). In paired measurements on dendrites and axons of the same cells, we observed that the diffusion coefficient in the axons was significantly higher than on the dendrites (D_axon_ = 0.29 ± 0.05 μm^2^/s, D_dendrite_ = 0.19 ± 0.05 μm^2^/s, mean ± s.e.m., *n* = 8 cells, *p* = 0.018, paired *t*-test).

The faster tracer diffusion on axons indicates fewer obstacles in the axonal membrane.^37^ Since the arrangement and area fraction of obstacles are not precisely known in the dendrites or axon, we cannot directly connect the difference in diffusion coefficients to a difference in membrane flow resistance. Nonetheless, an estimate with plausible parameters is informative: in a 2-dimensional fluid membrane, with randomly distributed MCA obstacles at an area-fraction of 0.1, a 2-fold difference in molecular diffusivity corresponds to a ∼20-fold difference in resistance to bulk membrane flow (Supplementary Calculation).^11,33^ Non-random obstacle arrangements could substantially amplify this difference.

A further indication of the unusual flow properties of axon membranes came from measurements of tether sliding. We pulled short tethers from axons and then translated the pipette parallel to the axon axis (Methods). In 14 of 23 tethers, the tether slid along the axon until it was perpendicular to the axon axis, minimizing the tether length (Fig. S2, Supplementary Movie 3). Similar tether sliding has been observed in axons of chicken sensory neurons^21^ and in axon terminals of goldfish retinal bipolar neurons.^26^ In contrast, tethers pulled from the soma remained pinned at their initial spots (Fig. S2), and others reported tether pinning in neutrophils,^22^ HeLa cells^21^, and adrenal chromaffin cells.^26^ This finding suggests that the spacing between MCAs in the axon exceeds the tether diameter (estimated to be 50 - 100 nm, Methods), whereas in non-motile cells the spacing between MCAs was estimated to be approximately 4 - 5 nm.^11^

### Membrane tension coordinates axon growth and branching

Growth and branching of the axon are required for the axon to navigate through the mechanically heterogeneous brain and form a properly connected network.^2^ Actin filaments push on the growth cone membrane to elongate the axon, and actin filaments deform the membrane on the axon shaft to initiate new branches. Could membrane tension play a role in coordinating the dynamics of growth and branching?

We hypothesized that a decrease in tension at the growth cone, triggered, for example, by a mechanical barrier which blocked growth cone extension, could propagate upstream along the axon and facilitate collateral branching. We used localized perfusion to lower tension at the growth cone and monitored upstream tension and branching.

First, in a neuron expressing cytosolic eGFP we pulled a tether 75 μm upstream of the growth cone and locally perfused the growth cone with deoxycholate (500 µM, Methods), which has been shown to lower local membrane tension.^38^ Application of deoxycholate at the growth cone led to a 50% drop in membrane tension at the upstream tether (Fig. 3A, 3B).

**Figure 3:**
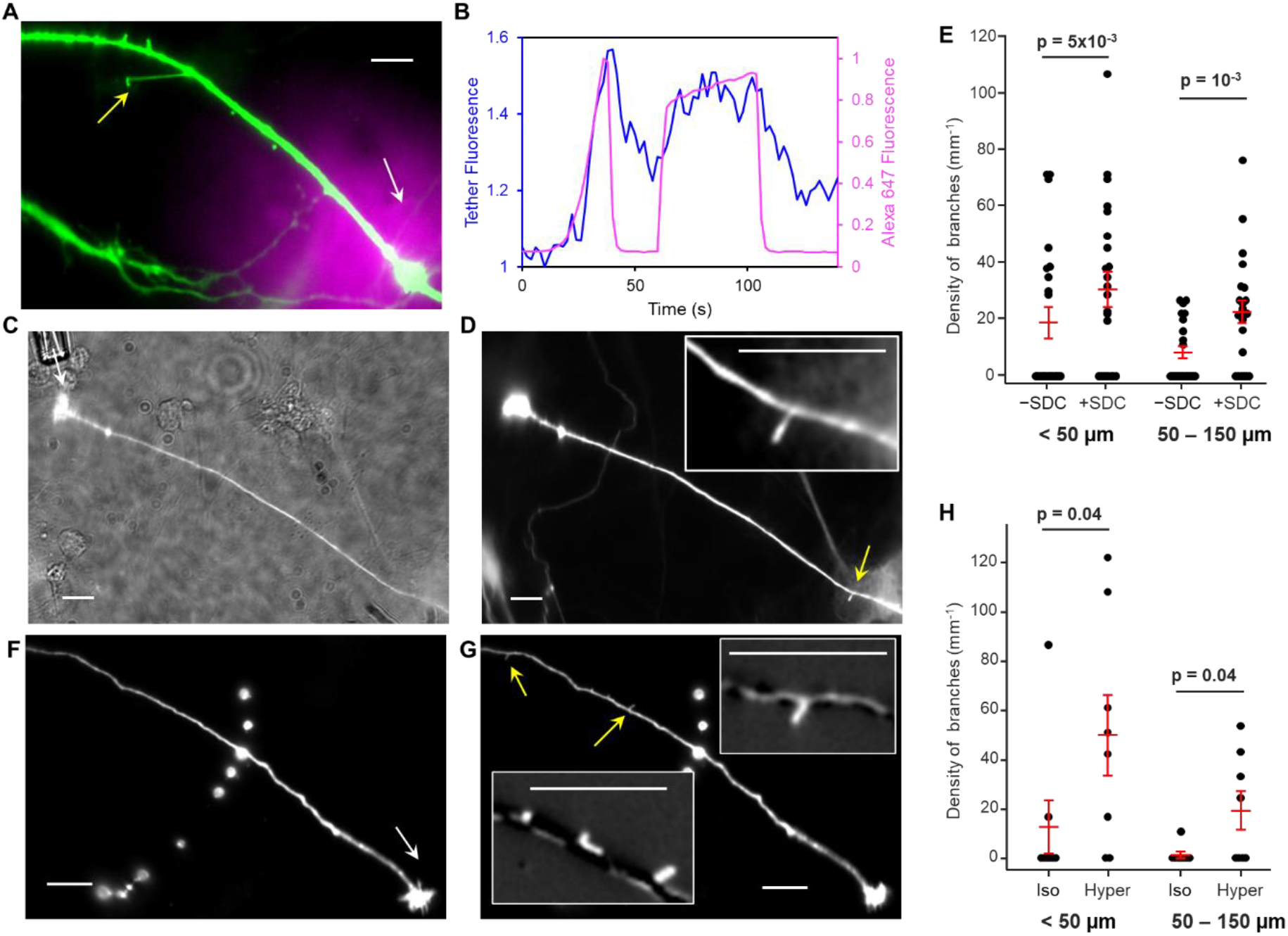
Lowering membrane tension at the growth cone triggers upstream collateral branching. **A)** Composite image showing local perfusion of deoxycholate, traced by Alexa 647 (magenta) and the growth cone (green) in a neuron expressing cytosolic eGFP. White arrow: perfusion pipette. Yellow arrow: membrane tether for tension sensing. **B)** Changes in tether fluorescence, inversely proportional to membrane tension (blue), in response to deoxycholate perfusion of the growth cone (magenta). **C**) Fluorescence and transmitted-light image of a neuron, with a deoxycholate-loaded pipette next to the growth cone (arrow). **D)** Perfusion of the growth cone with deoxycholate triggered formation of a new branch (arrow) in a distant section of the axon. Inset: close-up view of the new branch. **E)** Perfusion of deoxycholate (+SDC) at the growth cone increased the density of branches along the axon compared to before injection (-SDC). The same trend was observed close to (< 50 µm) to and far from (50 – 150 µm) the growth cone (n = 23 neurons). **F)** Fluorescence image of an axon before perfusion with hypertonic buffer. Arrow indicates the perfusion pipette. **G)** Hypertonic perfusion at the growth cone triggered collateral branching (arrows) in distant sections of the axon. Insets show magnified views of the change in fluorescence (after – before perfusion) highlighting new branches. **H)** Perfusion of hypertonic buffer (Hyper) at the growth cone increased the density of branches along the axon compared to before perfusion (Iso). The same trend was observed close to (< 50 µm) to and far from (50 – 150 µm) the growth cone (n = 8 neurons). P-values calculated from two-tailed paired *t* test. Scale bars 10 µm.

Local perfusion of the growth cone with deoxycholate also triggered initiation of new axonal branches outside the perfused region in 12/23 neurons (Fig. 3C, D, S3). The number density of putative branches along the axon significantly increased upon deoxycholate perfusion of the growth cone, with similar effect on axon segments close to the growth cone (< 50 µm, branch density from 19 ± 5 to 30 ± 6 mm^-1^, mean ± s.d., *p* = 0.005) and far from the growth cone (50 – 150 µm, branch density from 8 ± 2 to 22 ± 4 mm^-1^, *p* = 0.001), (Fig. 3E). Similar increases in branch density were observed when we injected hypertonic buffer to the growth cone (+350 mM mannitol, Methods), which reduces membrane tension through osmosis (Fig. 3F-3H).^39^ These experiments confirmed our hypothesis that lowering membrane tension at the growth cone increased axon collateral branching upstream.

Next, we asked whether an increase in axonal membrane tension, such as might occur due to new branching events, could inhibit extension of the growth cone or of other branches. To address this question, we pulled tethers on the axon at distances 25 to 110 μm from the growth cone, and monitored the motion of the growth cone and of other branches (Fig. 4A). The size of the growth cone decreased during tether pulling and recovered after relaxing the tether (Fig. 4B, 4C). Short branches near the growth cone also disappeared during the pulling period (Fig. 4C). On average, the growth cone shrank to 95.3 ± 1.3% of its original size during tether pulling and fully recovered (100.5 ± 1.3%) upon releasing the tether tension (mean ± s.e.m., *n* = 14 neurons, Fig. 4D). When we pulled a tether 110 µm from the growth cone, a clear 6 s delay in the growth cone response was observed (Fig. S4), consistent with our earlier measurements of membrane tension propagation speed ∼20 µm/s. The density of branches along the axon also decreased during tether pulling, from 22 ± 3 mm^-1^ to 11 ± 3 mm^-1^ (mean ± s.d., *p* = 0.003). The density of branches partially recovered to 17 ± 3 mm^-1^ within 1 min of releasing the tether (Fig. 4E).

**Figure 4:**
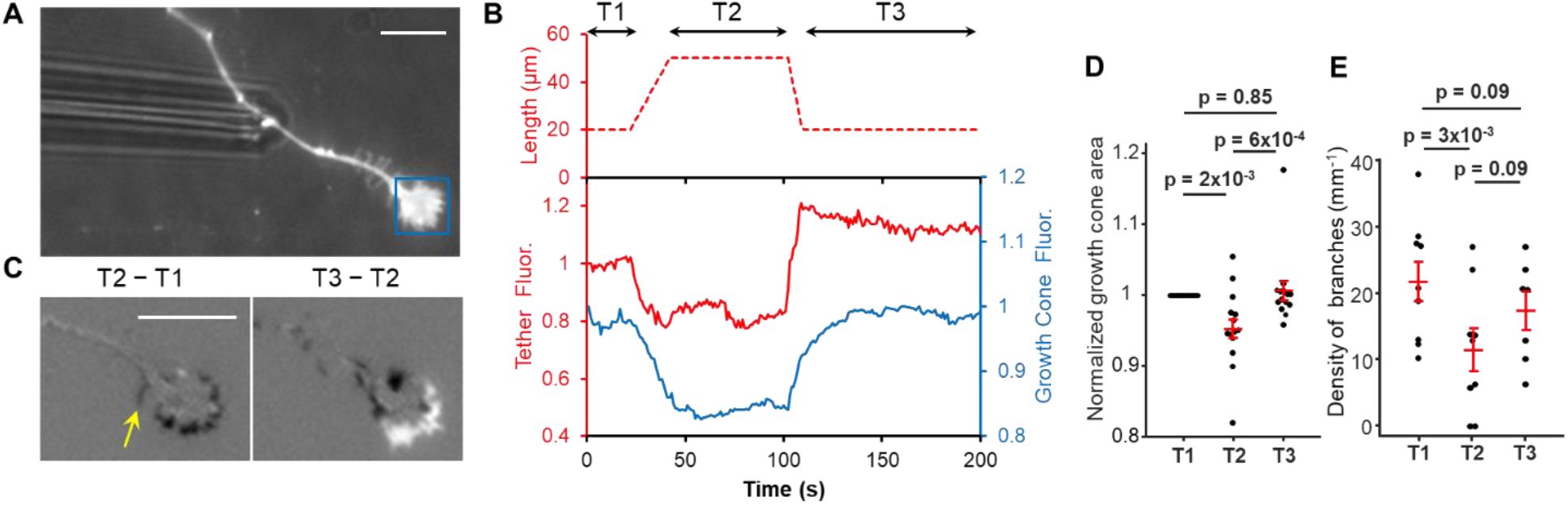
Increases of axonal membrane tension lead to retraction of growth cone and of collateral branches. **A)** Fluorescence and transmitted light image of a pipette holding a bead in contact with an axon expressing SEP-NXN. Blue box: growth cone. **B**: Top: tether length. Time windows show before (T1), during (T2), and after (T3) stretching. Bottom: fluorescence of the tether (red) and the growth cone (blue). **C)** Left: change in fluorescence after stretching the tether. Arrow points to the disappearance of one branch. Right: change in fluorescence after relaxing the tether. **D)** The growth cone shrank during tether pulling (n = 16 neurons) and re-grew after tether relaxation (n = 14 neurons). **E)** Number of branches on the axon decreased during tether pulling (n = 9) and partially recovered after tether relaxation (n = 7). In **E**, only axons with existing branches were chosen for tether pulling. P-values calculated from two-tailed paired t-test (in **D** the test was done on actual growth cone areas instead of the normalized areas), and n values refer to number of neurons. Scale bars 10 µm.

## Discussion

Our experiments show that changes in tension at the growth cone modulate branching rates, and that artificial induction of branches along the axon can trigger growth cone retraction. Together, these results suggest a role for long-range membrane tension signaling in axon pathfinding, shown schematically in Fig. 5. This picture is reminiscent of the model of neutrophil migration proposed by Houk and coworkers,^14^ in which tension propagation ensured that migration occurred along a single leading edge growth front. The key difference is that axons do not always migrate in a single direction, but sometimes contain branches. The axon trailing the growth cone leads to a gradual decoupling of tension over a ∼30 μm length-scale (Fig. 1E), permitting branches to nucleate if far enough apart, but maintaining a competition between extension of the growth cone and of nearby branches.

**Figure 5.**
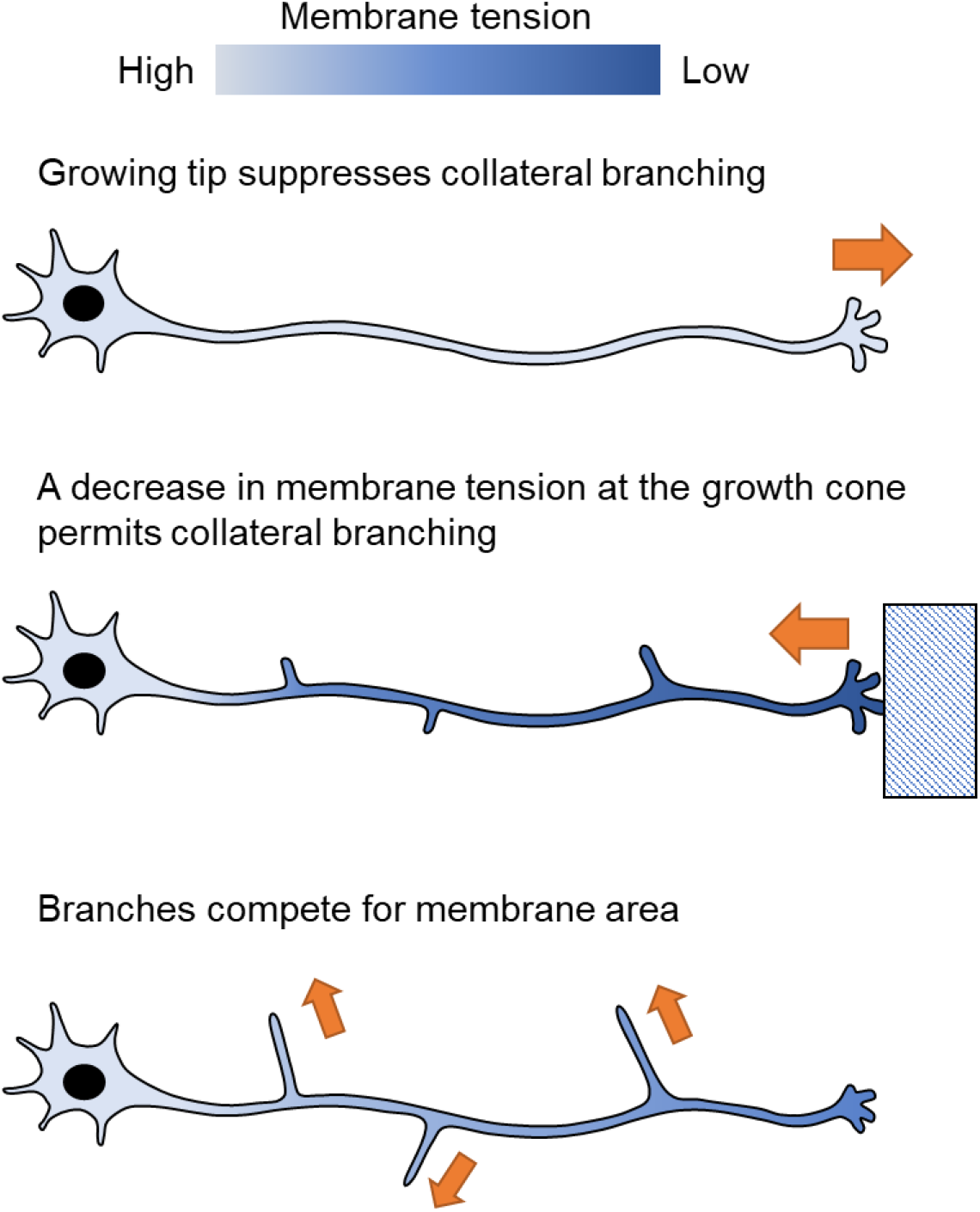
Model of membrane tension coordinating axonal growth and branching. During rapid growth cone advancement, membrane tension suppresses branching. Obstacles or mechanical heterogeneities that impede growth cone progress create a local imbalance between membrane addition and area expansion, leading to a propagating front of lower tension which enhances nucleation and growth of collaterals. As the collaterals grow, each broadcasts information about its progress to the others via long-range changes in membrane tension.

In prior studies on HeLa cells, a local perturbation to membrane tension was not detectable 5 - 10 μm away after a 10-minute delay,^10^ whereas here we showed that a similarly produced tension perturbation in axons propagated at ∼20 μm/s over tens of microns, a >1,000-fold difference in propagation speed. A critical challenge for future research will be to understand how and why this biologically important parameter varies across cell types and conditions, and how tension propagation couples physiological processes over long distances.

In a study on chick sensory axons, Dai and Sheetz reported a spontaneous retrograde membrane flow with a typical speed of 7 µm/min.^20^ This membrane flow speed is more than 100-fold slower than what we measured for tension propagation (20 µm/s). It is important to distinguish these two physical quantities: the speed of membrane flow is determined by the rate of mass transport and is linearly proportional to the gradient of membrane tension. In contrast, tension is expected to propagate diffusively^10^, so the effective propagation velocity depends on the measurement timescale and the stretching modulus of the membrane, but is expected to be approximately independent of the gradient in membrane tension.

Dai and Sheetz ascribed the retrograde membrane flow to a tension differential between the soma and the growth cone of ∼1.5 pN/µm. The lower tension at the growth cone was attributed to excess lipid deposited via fusion of trafficking vesicles. By combining the tension differential (1.5 pN/µm), the flow speed (7 µm/min) and the axon length (60 - 90 µm in ^20^), one obtains a membrane drag coefficient of ∼0.2 pN s/µm^3^ along the axon. In comparison, the drag coefficient in HeLa cells was reported to be ∼1600 pN s/µm^3^.^11^ The nearly 4 orders of magnitude difference in drag coefficient is consistent with the dramatically faster propagation of membrane tension in axons compared to HeLa cells.

The drag coefficient is proportional to the intrinsic lipid viscosity, which is not expected to vary substantially between cells, and inversely proportional to the Darcy permeability of the network of MCAs. Randomly distributed MCAs can cause substantial resistance to membrane flow in non-motile cells.^10,12^ However, it is possible to reduce the drag by simple rearrangements of MCAs, such as by clustering or aligning the anchored proteins.^11^ In axons, super-resolution imaging has revealed co-axial actin rings underlying the axonal membrane with 180 nm periodicity.^40^ The actin rings are linked by spectrin tetramers which bind to membrane proteins through ankyrin, resulting in highly aligned MCAs, that pose minimal resistance to axial membrane flow.^41,42^ Increased membrane flow may be a consequence of the periodic cytoskeleton in axons leading to clustering of MCAs. The tether sliding experiments (Fig. S2) confirm that MCAs are far more widely spaced in axons than in other cellular structures.

Membrane tension is an important regulator in cell migration, mechanosensing, and intracellular vesicle trafficking. Synaptic vesicle fusion adds lipids to, and lowers the membrane tension of, the presynaptic plasma membrane. Watanabe and coworkers, proposed that this tension drop can quickly travel across the active zone, triggering a mode of ultrafast endocytosis within 50 ms.^25^ Assuming a 500 nm size for the active zone, the speed of tension propagation must exceed 10 µm/s for this effect to occur, consistent with our estimates of tension propagation speed. While our study focused on the role of tension propagation in axon growth and branching, this does not rule out the possibility that tension propagation is involved in other axonal processes such as coordinating vesicle turnover.^26^

**Figure S1:**
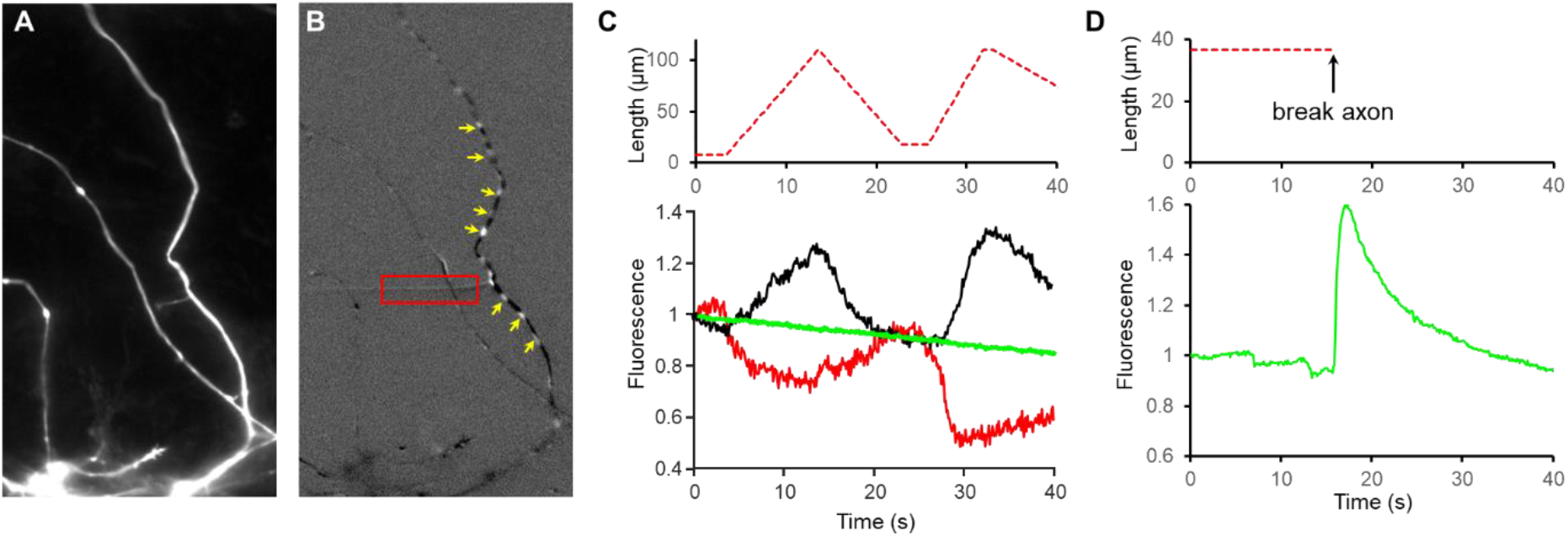
Propagation of membrane tension and axonal pearling are independent of Ca^2+^ signaling. **A)** In a rat hippocampal neuron expressing axon-targeted GCaMP6s, a membrane tether was alternately pulled and relaxed. **B)** Image of fluorescence changes when the tether was extended. The tether is in the red box, the pearls are indicated by yellow arrows. **C)** Fluorescence averaged along the length of the axon (green) was constant except for slow photobleaching. Fluorescence of the tether (red), and average of the pearls (black) showed opposite responses to tether stretch (red dash). **D)** Breaking the axon in **A** with a pipette at t = 16 s evoked a large Ca^2+^ transient signal. Example is a 10 DIV neuron, representative of experiments on *n* = 5 neurons (7 - 14 DIV).

**Figure S2:**
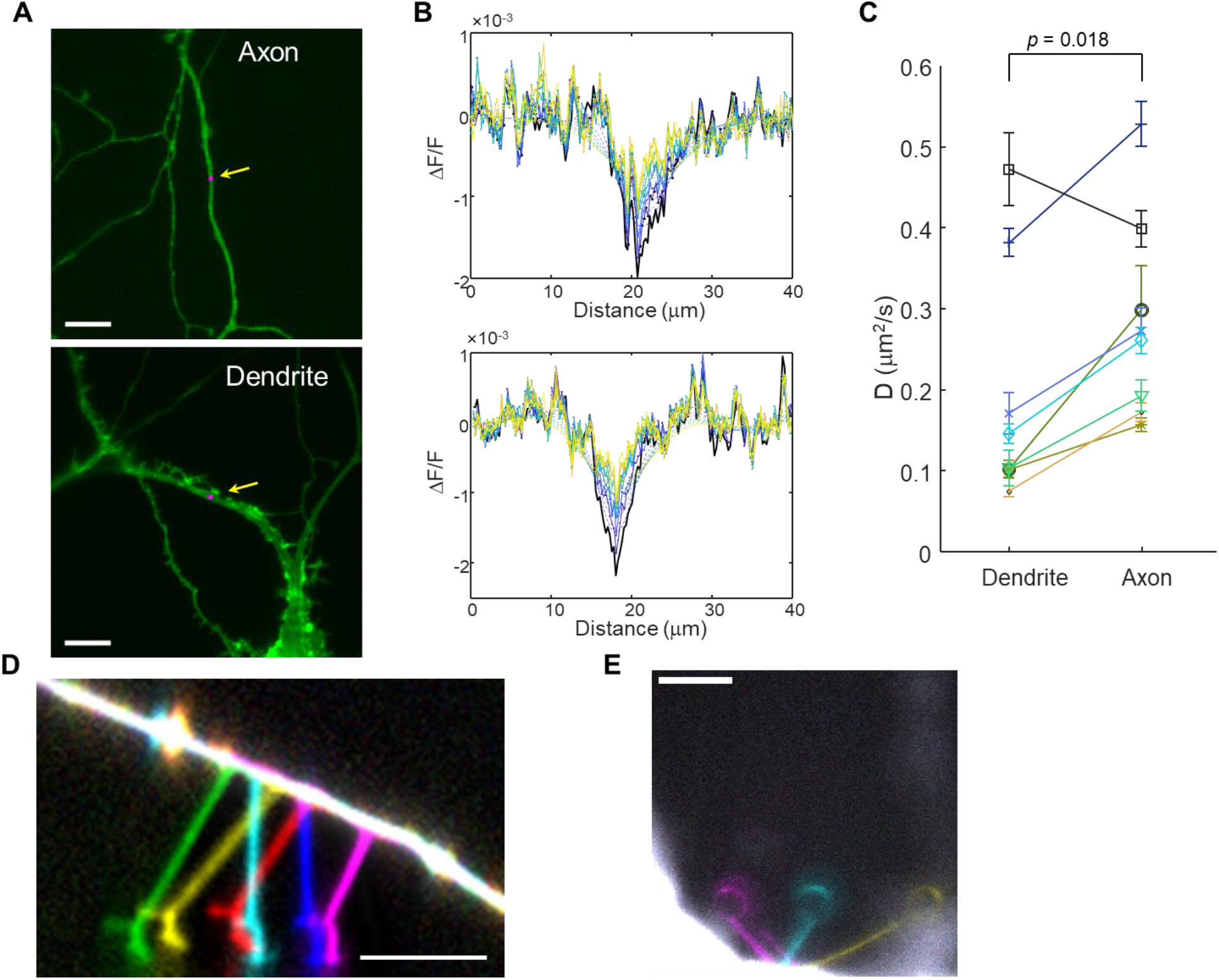
Axon membranes have fewer obstacles than dendrite membranes. **A)** Composite images showing the neuron expressing GPI-eGFP (green) and a 1 µm diameter photobleaching spot (magenta) on the axon (top), and dendrite (bottom). Scale bars 10 μm. **B)** Fitting of the FRAP data. Line profiles through the photobleached spot were measured every 5 s for 50 s. The profile at each time was fit by convolving the initial profile with a Gaussian diffusion kernel. The diffusion coefficient was varied to optimize the global fit. **C)** Pairwise measurements of GPI-eGFP tracer diffusion from dendrite and axon. The diffusion coefficient was 78 ± 62% higher in the axons than the dendrites (mean ± s.d., *n* = 8 neurons). **D)** Sliding of a tether along the axon (left then right). Colors: red, yellow, green, cyan, blue, magenta correspond to images taken at time 0, 2, 5, 9, 11, and 15 s respectively. **E)** Tether pinning on a neuron soma. Colors magenta, cyan, yellow correspond to images taken at 1, 80 and 100 s. Scale bars 10 µm.

**Figure S3:**
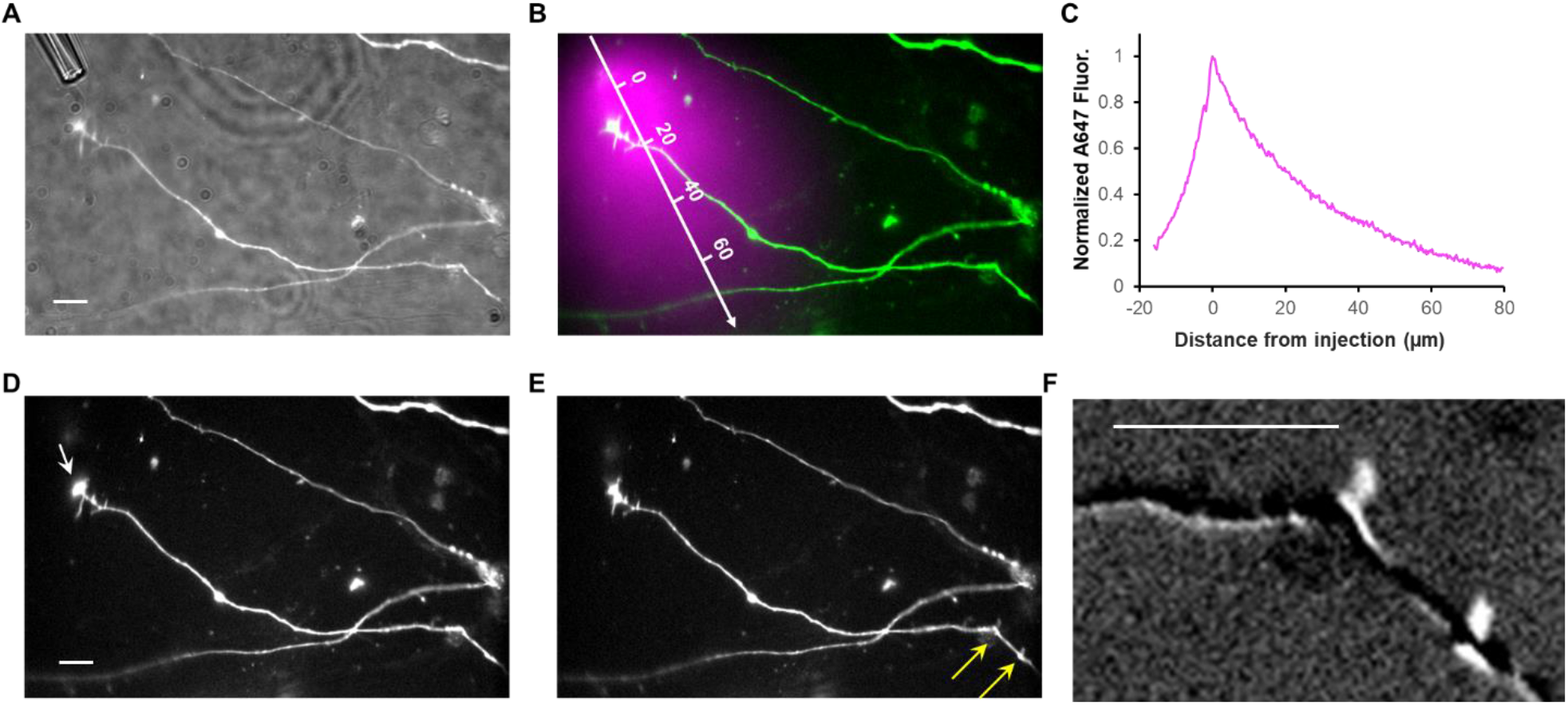
Injection of deoxycholate at the growth cone triggers upstream axon branching. **A)** Composite (fluorescence and transmitted light) image of a neuron expressing SEP-NXN, with a deoxycholate-loaded injection pipette next to the growth cone. **B)** Composite fluorescence image showing the perfusate profile (traced via Alexa-647, magenta) relative to the axon (SEP-NXN, green). **C)** Line profile of Alexa-647 fluorescence along the axis shown in (B). **D)** Axon before deoxycholate injection (arrow indicates the injection pipette). **E)** Formation of new branches (arrows) within 1 minute after perfusing deoxycholate at the growth cone. **F)** Close-up view of the new branches showing fluorescence after perfusion minus fluorescence before perfusion. Scale bars: 10 µm.

**Figure S4:**
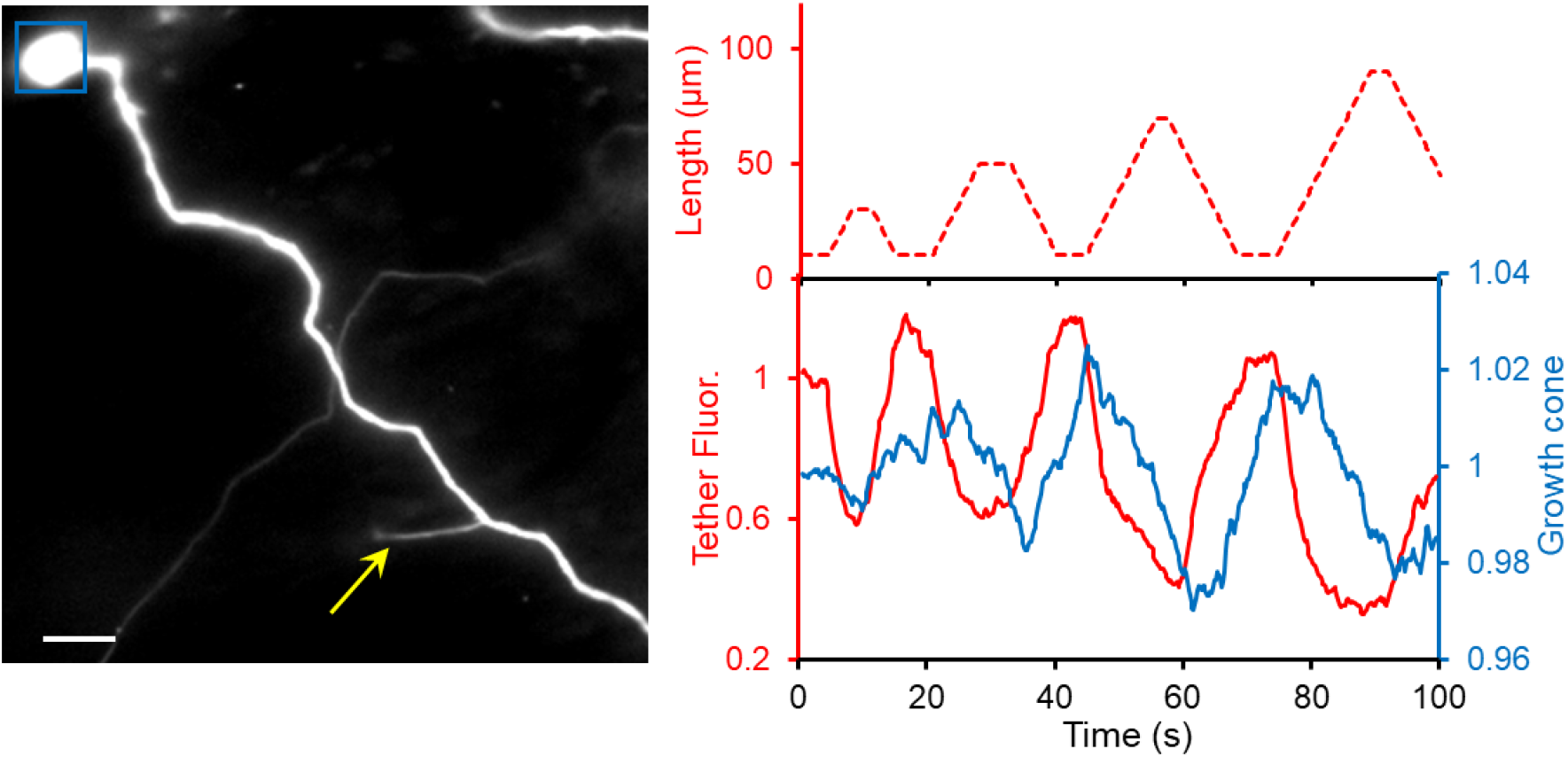
Long-range coupling between membrane tension and growth cone size. Left: fluorescence image showing an axon expressing SEP-NXN, with a tether pulled 110 µm upstream from the growth cone (arrow). Right: Tether length (top) and relative changes of the tether fluorescence (red) and growth cone area (blue), showing a 6 s delay of the growth cone response relative to the tether response. Scale bar 10 µm.

## Supporting information

Movie S1: pulling two tethers from an axon

Movie S2: pulling a tether can induce pearling

Movie S3: tether sliding along an axon

## Supplementary Calculation: Effect of MCA density on tracer diffusion and membrane flow

The density of immobile obstacles affects both the diffusion of tracers and the flow of membrane. Let *ϕ* be the area fraction of membrane obstacles. We previously showed that at low *ϕ*, the tracer diffusion coefficient, *D*_T_ scales as 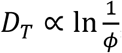, while the Darcy permeability scales as 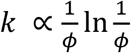.^11^ Consider two cellular compartments that differ only in *ϕ*, with tracer diffusion coefficients *D*_1_ and *D*_2_. If the ratio *D*_2_/*D*_1_ = *α*, then rearrangement of the above proportionalities yields:

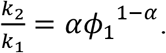

For example, if *α* = 2 (i.e. the diffusion coefficients differ by a factor of two) and *ϕ*_1_ = 0.1, then *k*_*2*_/*k*_1_ = 20. If *α* = 4 and *ϕ*_1_ = 0.1, then *k*_*2*_/*k*_1_ = 4000. This calculation assumes a random distribution of homogeneous disk-like obstacles. Clustering of obstacles in region 2 could further enhance the ratio *k*_2_/*k*_1_.

### Supplementary movie captions

Supplementary Movie 1: Pulling two tethers 40 µm apart on an axon. The amplitude of the fluorescence changes in tether 2 was 0.33 of the fluorescence changes in the pulled tether, tether 1. Tether pulling also induced pearl formation in the axon.

**Supplementary Movie 2:** Tension-induced axon pearling. The baseline fluorescence has been subtracted from the movie to highlight the changes in fluorescence due to axonal pearling.

**Supplementary Movie 3:** Tether sliding along and axon. Movie is shown at 5x real-time.

## Acknowledgements

We thank Irina Shlosman and Shahinoor Begum for technical assistance. We thank Kranthi Mandadapu, Ahmed El Hady, Erdem Karatekin, and Ruobo Zhou for helpful discussions. This work was supported by the Howard Hughes Medical Institute, a Vannevar Bush Faculty Fellowship, and a Jane Coffin Childs Fellowship Award to S.I.G.

## Methods

### Neuron culture

All procedures involving animals were in accordance with the National Institutes of Health Guide for the care and use of laboratory animals and were approved by the Institutional Animal Care and Use Committee (IACUC) at Harvard. Hippocampal neurons from P0 rat pups were purchased from BrainBits and cultured in NBActiv4 medium at a density of 5,000 to 30,000 cells/cm^2^ on glass-bottom dishes pre-coated with poly-d-lysine and Matrigel. At 1 day in vitro (DIV), glia cells were plated on top of the neurons at a density of 7000 cells/cm^2^. At 3 - 5 DIV neurons were transfected following the calcium phosphate protocol (Jiang and Chen, 2006). Imaging was performed on DIV 7 to 14, with neuron culture medium replaced with extracellular (XC) imaging buffer (125 mM NaCl, 2.5 mM KCl, 15 mM HEPES, 30 mM glucose, 1 mM MgCl_2_, 3 mM CaCl_2_, and pH 7.3).

### Neuron labeling

For experiments in Figs. 1, 2, S2, 3A-D, neurons were labeled with cytosolic eGFP. We found that a membrane tag comprised of superecliptic pHluorin-neurexin (SEP-NXN, construct ZS090) improved axonal membrane visualization, so for experiments in Figs. 3F-H, S3, 4, and S4, we labeled neurons with SEP-NXN.

### Tether pulling

Micropipettes were pulled from glass capillaries (World Precision Instrument, 1B150F-4) using a pipette puller (Sutter P1000). The tip of the pipette was cut to an opening diameter of ∼3 μm and bent to ∼40° using a microforge (WPI, DMF1000). Tethers were pulled with a 4 µm diameter polystyrene bead (Spherotech #DIGP-40-2) held at the tip of a micropipette and controlled by micromanipulators. In cases where tethers broke, the piece of tether attached to cells retracted in about 1 min, confirming that there were not cytoskeletal elements within the tethers.

### Cloning and constructs

SEP-NXN (ZS090): To facilitate the identification of growth cones, we switched the axon marker from cytosolic eGFP to the axon specific rat neurexin-1β (NXN), tagged with a super-ecliptic pHluorin (SEP) at the N-terminus (SEP-NXN). SEP from Addgene # 24000,^43^ NXN from Addgene #44968,^44^ were ligated together via Gibson cloning. GPI-eGFP was from Addgene #32601.^45^ Cytosolic eGFP was used for most of the tether imaging. Axon GCaMP6s (Addgene # 112005) was from Ref. ^32^.

### Perfusion experiments

A micropipette was loaded with XC buffer supplemented with either sodium deoxycholate (500 µM, Sigma 30970) or D-Mannitol (350 mM, Sigma M4125). The micropipette was then positioned ∼10 µm away from the growth cone (see Fig. S3). A slight suction pressure was used to minimize the leakage of the solution inside the pipette before experiments. While imaging of the axon, a pressure ∼0.1 atm was used to inject the solution inside the pipette for 10 - 30 s, aiming at the growth cone. The axon was continuously monitored until > 1 min after stopping the injection.

### Estimate of tether diameter

When tethers were pulled from an axon expressing fluorescent membrane markers, tether diameters were estimated by comparing the background-corrected cumulative cross-sectional fluorescence of the tether vs. axon. The integrated fluorescence is proportional to the surface area, which is proportional to the radius.

### Estimate of diffusion coefficients

Localized photobleaching was performed using a 488 nm laser focused to a small spot on the face of a digital micromirror device (DMD). The DMD was used to restrict the illuminated spot to a 1 μm circle aligned on the axon or dendrite of a rat hippocampal neuron expressing GPI-eGFP. The spot was bleached for 15 s and recovery was monitored for 5 min, with most of the recovery occurring within the first ∼30 s. The fluorescence intensity profile *F*(*r,t*) was extracted along the profile of the axon or dendrite and the first 50 s of recovery were fit to a 1D diffusion model:

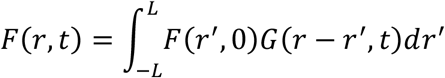

where the Gaussian diffusion kernel is:

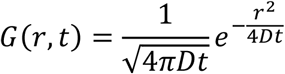

with *D* as the only free parameter. Image analysis and nonlinear least squares fitting were carried out in MATLAB.

## Notes

### Competing Interest Statement

The authors have declared no competing interest.

